# Ancient Reconstructed Proteins: A Framework for Resurrecting Protein Structures from Million-Year-Old Metagenomes

**DOI:** 10.64898/2026.08.20.745946

**Authors:** Louis Kraft, Peter Wad Sackett, Gabriel Renaud

## Abstract

DNA sequences derived from ancient samples provide insights into human history, paleoenvironments, and evolutionary biology. Advances in laboratory techniques and computational tools have established ancient DNA research as a distinct field. However, current analyses focus mainly on the DNA level, while the protein space remains underexplored. Recent progress in the *de novo* assembly of ancient metagenomes and the availability of protein structure prediction tools, such as AlphaFold 2, enable the reconstruction of protein structures from these degraded sequences. Here, we present a computational framework to assemble contigs, evaluate their authenticity as ancient sequences, predict open reading frames, and fold ancient protein structures directly from highly damaged metagenomic data. Applying this pipeline to two-million-year-old datasets from the Kap København Formation, we successfully rescued ancient proteins involved in methane metabolism. By generating structural models with AlphaFold 2 and comparing them to modern predicted reference structures, we demonstrate that these ancient proteins can be reconstructed and aligned with high confidence. We showcase this by analyzing an archaeal V/A-type ATP synthase protein recovered from the 2M-year-old Greenlandic data. Ultimately, our work proves that ancient proteins can be reliably recovered from highly degraded palaeogenomic material, establishing a new computational avenue for evolutionary and biochemical research.

## 1 Introduction

Recent advances in protein structure prediction have transformed protein biology. Methods such as AlphaFold 2 enable the accurate inference of three-dimensional protein structures directly from amino acid sequences [1], leading to the expansion of databases such as the AlphaFold database, which now contains over 214 million predicted structures [2]. Combined with over 595 million sequences in the NCBI nr database [3], these resources represent a vast knowledge base for functional and evolutionary research [4]. Structure-based comparison tools now facilitate large-scale functional screening, allowing researchers to place protein sequences in a structural context that extends beyond sequence similarity alone [5, 6, 7]. However, these breakthrough technologies have not yet been systematically applied to ancient metagenomics, even though extensive reservoirs of ancient protein-coding sequences exist within publicly available datasets which could provide valuable insights into the evolution of structure and function [8].

Ancient metagenomics has evolved from reconstructing individual pathogens in host-associated samples to characterizing entire paleoecosystems from environmental sediments and permafrost [9, 3]. This expansion enables the study of past microbial dynamics and biogeochemical cycles, providing insights into evolutionary history and historical climate responses [10, 11]. Despite these successes, research remains predominantly focused on the nucleotide level, utilizing taxonomic profiling and gene-centric searches to infer the functional potential of ancient organisms.

Translating aDNA into reliable protein structure models poses specific challenges that have hindered this integration. Ancient DNA molecules are highly fragmented, often resulting in sequencing reads of 50bp or less [12]. Additionally, post-mortem cytosine deamination produces characteristic C*→*T and G*→*A substitutions at read termini [13, 14]. Furthermore, ancient metagenomic samples contain DNA from numerous taxa with uneven abundances, increasing the risk of chimeric assemblies and modern contamination [11, 15, 16]. These properties complicate *de novo* assembly and downstream gene prediction.

Despite these hurdles, recent work has demonstrated the potential of ancient assemblies for functional discovery. For example, Klapper *et al.* [17] used *de novo* assembly of ancient metagenomes to identify previously unknown biosynthetic gene clusters or Hodgins *et al.* [18] who reconstructed *in vivo* an ancient neurotoxin from *Clostridium tetani*. While this illustrates that ancient sequence data can provide access to unexplored functional diversity, such analyses remain strictly sequence-based. Expanding these approaches to structural modelling requires highly stringent validation. Specifically, it requires computational tools capable of rapidly and accurately evaluating ancient damage patterns across large-scale metagenomic datasets to authenticate the assembled open reading frames (ORFs) before the computationally intensive protein folding is done.

Here, we present a workflow that bridges these domains: from ultra-short, damaged ancient DNA reads to reconstructed protein sequences and their three-dimensional structural models. The pipeline which combines recently published methods integrates damage-aware assembly, gene prediction, rapid authentication of ancient reads, and structure prediction using state-of-the-art tools. To demonstrate its feasibility, we applied this workflow to highly diverse datasets from the *≈* 2.5-million-year-old Kap København Formation in North Greenland [9]. Guided by previous findings of enriched methanogenesis genes [10], we focused on reconstructing proteins involved in methane cycling. To illustrate the potential of our approach, we reconstructed an archaeal V/A-type ATP synthase subunit A and analyzed the resulting fold. The predicted fold was essentially indistinguishable from modern variants while offering amino acid substitutions absent from modern sequences. By incorporating structural information, our approach extends aDNA analysis beyond the nucleotide level and enables the functional characterization of ancient proteins in a three-dimensional context, providing a template for future protein-centric palaeogenomic investigations.

## 2 Materials and Methods

For this study, we employed a series of tools and workflows to process and analyse the Kap København metagenomic data. An overview of the procedure, including the main tools used in each step, is shown in Figure **1**.

**Figure 1:**
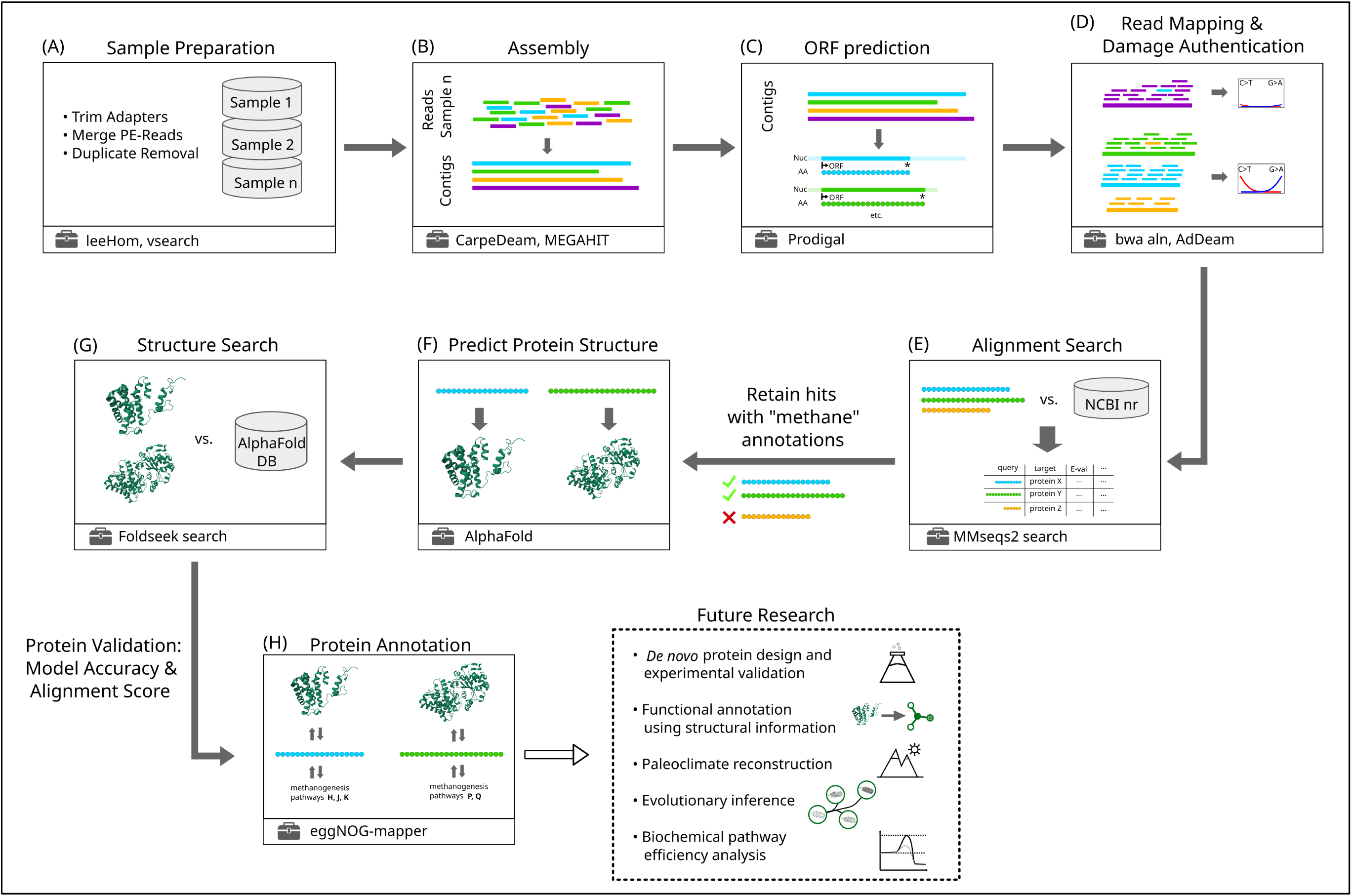
Workflow overview: (A) raw aDNA read retrieval and pre-processing; (B) metagenomic assembly; (C) ORF prediction; (D) aDNA authentication; (E) database search and methane-related ORF filtering; (F) protein structure prediction; (G) structural clustering and database search; (H) functional annotation.

### 2.1 Data Preparation

This step is shown in Figure **1**, **panel A**. The raw reads of the Kap København data were retrieved from the European Nucleotide Archive (ENA) under project number PRJEB55522. We downloaded both forward and reverse reads for a total of 67 different samples originating from various sampling locations, as described in the original studies [9, 10].

Prior to merging, the raw reads were processed with seqkit sana [19] to remove unpaired and corrupted reads. The sanitized reads were then processed with leeHom [20] for adapter trimming and paired-end merging with the --ancientdna flag. To remove PCR duplicates and retain only reads of length at least 30 bp, we used vsearch [21] with the flags --strand both and --minseqlength 30.

Basic statistics for these processed reads, collectively representing all 67 datasets, are shown in Table 1.

**Table 1:** Summary statistics for DNA sequences in FASTQ format as reported by seqkit stats [19].

| # reads | sum len | min len | max len | N50 | AvgQual | GC(%) |
| --- | --- | --- | --- | --- | --- | --- |
| 9,259,031,240 | 525,201,711,977 | 30 | 191 | 59 | 38.48 | 46.31 |

### 2.2 Assembly

All reads were assembled using two different assemblers. First, we ran MEGAHIT [22] with default parameters. Second, we ran CarpeDeam [15] in both safe and unsafe modes, also with default parameters except for the chosen mode. Shown in **panel B** of Figure **1**.

After assembly, Prodigal [23] was used in meta mode to identify and extract open reading frames (ORFs). Only complete ORFs (those containing both a start and a stop codon) were retained for downstream processing (Figure **1**, **panel C**).

### 2.3 aDNA Authentication

To assess aDNA authenticity, the processed reads were mapped back to the assembled contigs using bwa aln [24] with parameters -l 1024 -o 2 -n 0.01. The SAM files resulting from this mapping were converted to BAM files with samtools [25], and MD-flags were added to annotate matches and mismatches.

For each of the 67 samples, three BAM files (one per assembler) were generated. These were processed with AdDeam [26] in meta mode to generate a damage profile for each ORF (in nucleotide space) and cluster ORFs with similar DNA damage patterns. By default, AdDeam clusters patterns for various values of *k*; we focused on *k* = 3 to distinguish no damage, moderate damaged, and high damage. Contigs from any cluster with at least a 5% C*→*T substitution rate in any of the first five positions were retained for downstream analyses. This step is illustrated in Figure **1**, **panel D**.

### 2.4 Database Search and Contig Filtering

We searched protein sequences predicted by Prodigal against the NCBI nr database [3] using MMseqs2 [27] with --search-type 1, running on a high-performance computing (HPC) cluster. Only protein sequences whose corresponding contigs (in nucleotide space) exhibited characteristic ancient DNA damage were included in the search. Protein sequences were not clustered prior to this search to avoid discarding potentially informative hits.

Over 720,000 “ancient” ORFs were queried, producing large BLAST-like result tables in TSV format. We first retained only one hit per database target (based on lowest E-value) and then merged the TSV outputs from the three assemblies of each sample, ultimately keeping only the best hit per target across all assemblies (also based on lowest E-value). Finally, we restricted our analysis to hits annotated with the term “methane” in the NCBI nr database description. This procedure considerably reduced the number of protein sequences to be explored in further analyses (Figure **1**, **panel E**). These filtering criteria were designed to yield a tractable set of candidate proteins for downstream structure prediction. A more sensitive filtering strategy or a larger-scale structure prediction effort could uncover additional candidate proteins, offering a natural extension for future work.

### 2.5 Protein Structure Prediction and Structure Search

After filtering by matching the term “methane” in the database annotations, we were left with 2,004 sequences putatively involved in methane-related pathways. To predict their structures, we used the localColabFold repository (https://github.com/YoshitakaMo/localcolabfold), which uses ColabFold [28] and AlphaFold [1]. We ran this workflow with the parameters --num-seeds 2 --num-model 5 --num-recycle 3, producing 10 protein structure models for each sequence, and selected the model with the highest pTM score for further analysis. This process is shown in Figure **1**, **panel F**.

Next, we clustered the resulting structures using Foldseek cluster [4] (-c 0.9 -e 0.01 --min-seq-id 0.3), following the criteria of Barrio-Hernandez *et al.*, ultimately obtaining 1,299 unique protein structures (the representative structures of the clusters). These structures were then searched against the AlphaFold database [2] using Foldseek [5], retaining only the best hit per database target (lowest E-value). See **panel G**, Figure **1**. The results are presented in Figure **3**, Section “Protein Structure Evaluation”. Because the AlphaFold database entries are not necessarily experimentally validated, we compared aDNA-derived structures to their respective AlphaFold matches by considering pLDDT values and alignment confidence (pTM scores).

### 2.6 Annotation

To gain insights into the metabolic pathways associated with the ancient protein structures, we employed eggNOG-mapper [29], which assigns orthology-based functional terms, including KEGG pathway identifiers [30], to our sequences (Figure **1**, **panel H**). The outputs were further analysed with the kegg-pathways-completeness-tool (https://github.com/EBI-Metagenomics/kegg-pathways-completeness-tool) to evaluate pathway completeness and identify potentially relevant metabolic functions.

## 3 Results

### 3.1 Sequence Assembly and Database Search

As described in Section 3.2, we assembled 67 ancient metagenomic samples using multiple assembler configurations to maximise sequence recovery. The advantage of using several assemblers was shown in previous research [15]. Specifically, we *de novo* assembled each sample with MEGAHIT [22] and the damage-aware assembler CarpeDeam in both safe and unsafe modes and retained only contigs longer than 500 bp. In total, we obtained over 7.7 million contigs, summing to more than 11.8 billion base pairs. At this stage, we did not perform redundancy reduction to ensure that the full spectrum of nucleotide diversity was preserved. Key assembly metrics such as number of contigs, cumulative length, and related statistics are summarised in Table 2. The N50 value is relatively low (2,054 bp), reflecting the challenge of assembling highly degraded ancient metagenomic datasets. However, since our objective was to recover protein-coding regions, these contig lengths were sufficient for our objectives including translation and structural modelling.

**Table 2:** Assembly statistics for contigs *≥*500 bp across 67 samples, showing total contig count, cumulative length, length range, N50, and GC content.

| # contigs | sum len | min len | avg len | max len | N50 | GC(%) |
| --- | --- | --- | --- | --- | --- | --- |
| 7,744,567 | 11,827,396,520 | 500 | 1,527.2 | 75,062 | 2,054 | 49.79% |

We extracted all full-length ORFs from the assembled contigs and remapped the original reads to these ORFs (see Section 3.3). Rather than plotting each ORF’s individual damage profile, we used the representative clusters inferred by AdDeam [26]. Thus, Figure **2**, **panel A**, displays for each ORF the substitution frequencies (C*→*T and G*→*A) at the first five and last five positions of the reads mapping to that ORF. Because each of the 67 samples was clustered into *k* = 3 damage profiles, there are 201 representative profiles in total, but we only show those with at least 5% C*→*T substitution in any of the first five positions. Most ORFs flagged as ancient by this criterion exhibit very high C*→*T substitution rates (*≥*40%) at the first nucleotide position of the mapped read. In total, we extracted 14,947,770 full-length ORFs from the contigs. After filtering out those that did not meet the damage threshold or had fewer than 100 reads mapped, 726,353 ORFs remained for further analysis.

To identify sequences that are potentially involved in methane metabolism, we searched all ancient ORFs in protein space against the NCBI nr database [3], which provides broad taxonomic coverage and a baseline of functional annotation. At this stage, we had curated for each sample three sets of presumably ancient ORFs, one from each assembler configuration. We then searched all ORFs (in amino acid space) against the database, producing large BLAST-like TSV files with multiple hits per query. These files included both full-length and partial alignments. Following the procedure in Section 3.4, we filtered the TSV files to retain only the single best hit per ORF and per target defined by the lowest E-value, thereby excluding any alternative hits produced by the other two assemblers.

**Figure 2:**
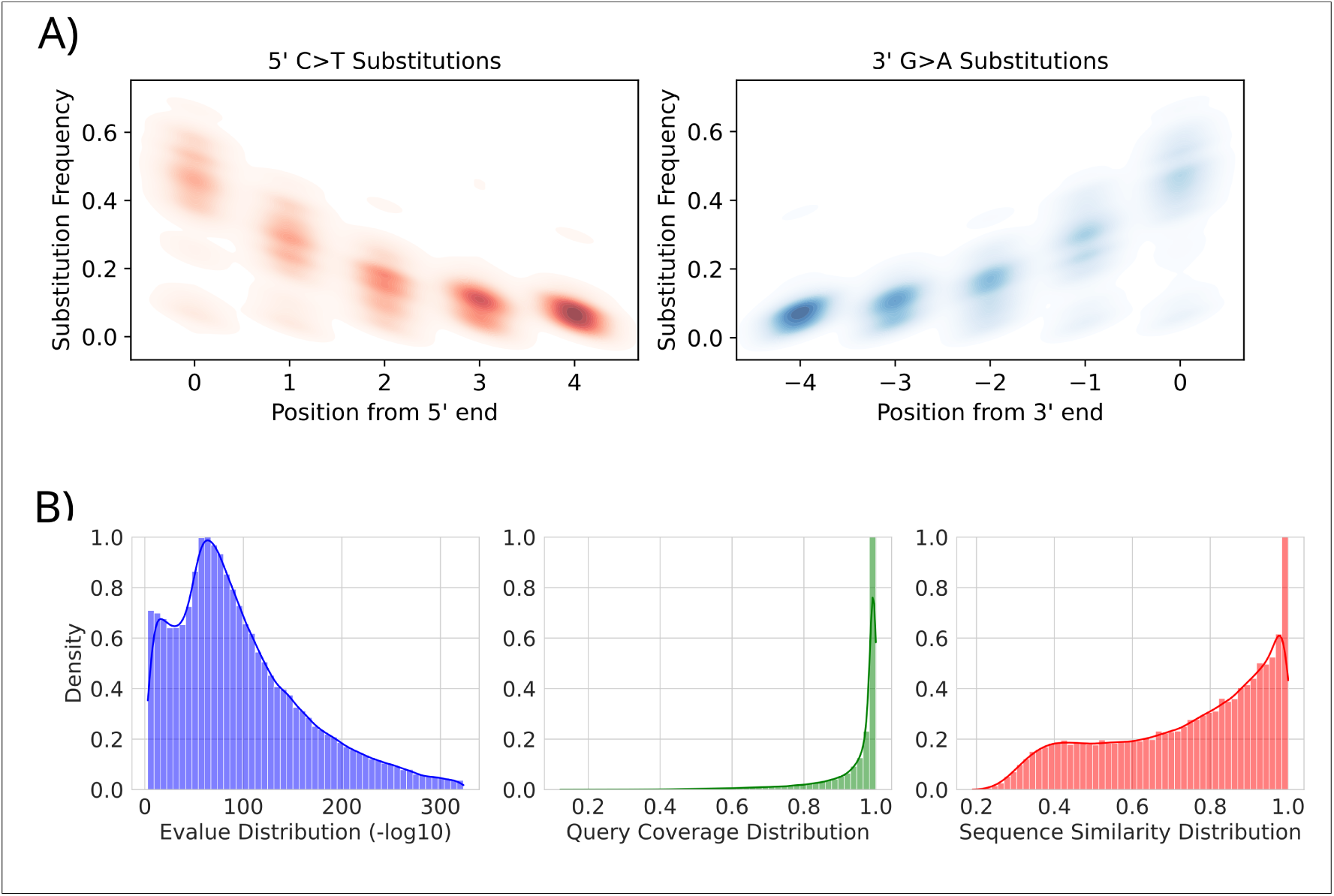
Damage patterns and sequence search metrics for ancient ORFs. **Panel A:** C*→*T and G*→*A substitution frequencies at the first and last five positions of reads mapping to ORFs with *≥*5% damage. Shown are 201 representative damage profiles as reported by AdDeam, rather than all 726,353 individual profiles. **Panel B:** Distributions of best-hit E-values, query coverage, and sequence similarity for filtered ORFs against the NCBI nr database.

Figure **2**, **panel B**, summarizes three metrics from the sequence search results: distribution of E-values, query coverage, and sequence similarity. The latter is defined as number of matches over the aligned length. All results refer to the best hits only, which is on hit per assembled and predicted ORF and one hit per target only. The E-value distribution (left) shows that the vast majority of protein hits fall between 1 and 10*^−100^*, with counts gradually declining at lower Evalues. A small fraction of hits achieve E-values below 10*^−300^*. The query coverage (middle) shows a pronounced peak at 100%, indicating that most ORFs align over their entire length. In contrast, sequence identity (right) is more broadly distributed: the number of hits rises almost linearly from 0–40% similarity, remains relatively uniform between 40–70%, and then increases towards 100%, with a distinct peak at 100% sequence similarity.

Notably, 76,482 predicted ORFs of supposedly ancient origin did not match with any sequence in the NCBI nr database with our search approach. This finding is explored further in the Discussion.

### 3.2 Protein Structure Evaluation

A goal of this study was to investigate the three-dimensional structures of the ancient proteins we extracted from the assembly. However, structure prediction is computationally intensive, requiring computation of multiple sequence alignments and GPUs for structure modelling. To narrow down the 726,353 candidate proteins, we kept only those whose top hit in the NCBI nr search was annotated with the substring “methan”, thereby selecting both methanotroph- and methanogen-related sequences and reducing the dataset to a computationally manageable size. Following the procedure in Section 3.5, we selected 2,004 protein sequences for structure prediction with AlphaFold 2 [1]. The resulting models were then clustered by structural similarity using the Foldseek cluster module [4]. After clustering and filtering, 1,299 representative structures remained: 542 from MEGAHIT assemblies [22], 427 from CarpeDeam safe mode [15], and 330 from CarpeDeam unsafe mode [15].

Finally, we compared our predicted ancient protein structures against the AlphaFold db [2] to assess their similarity to modern homologues. Because in silico models are not inherently accurate, we also examined confidence metrics for both our predictions and the database entries [31]. AlphaFold reports two key scores: the predicted Local Distance Difference Test (pLDDT), which reflects per-residue confidence (scale 0 to 100), and the predicted Template Modeling score (pTM), which assesses overall fold accuracy on a scale from 0 to 1 [1, 32]. These metrics were originally developed to compare models against experimentally determined structures [33, 34], but AlphaFold’s neural network predicts proxy values for them directly from internal sequence representations generated during modelling [1].

However, pTM scores were only available for our own predicted protein structure models, as AlphaFold db entries do not report the pTM score. In contrast, average pLDDT values could be obtained for both our ancient predictions and the corresponding AlphaFold db structures.

In Figure **3**, **panel A** (left), we plot each ancient model’s pTM score against its average pLDDT score, with points color-coded by the alignment TM score to the best hit in the AlphaFold db. pLDDT values above 90 are generally considered high-confidence predictions [31, 35]. Although there is no universally accepted pTM cutoff, a pTM score over 0.5 typically indicates overall fold similarity in traditional structural comparisons [36], and has been proposed for pTM evaluation [37]. For structural alignment assessment, TM scores above 0.8 indicate highly accurate structural matches [38, 39]. We colored data points red when their alignment *≥*0.8, thereby highlighting ancient protein models that closely match a modern reference structure in the database. 1,074 out of the 1,299 representative protein models had a model pLDDT *≥* 0.9 and aligned to references in the AlphaFold database with TM *≥* 0.8.

**Figure 3:**
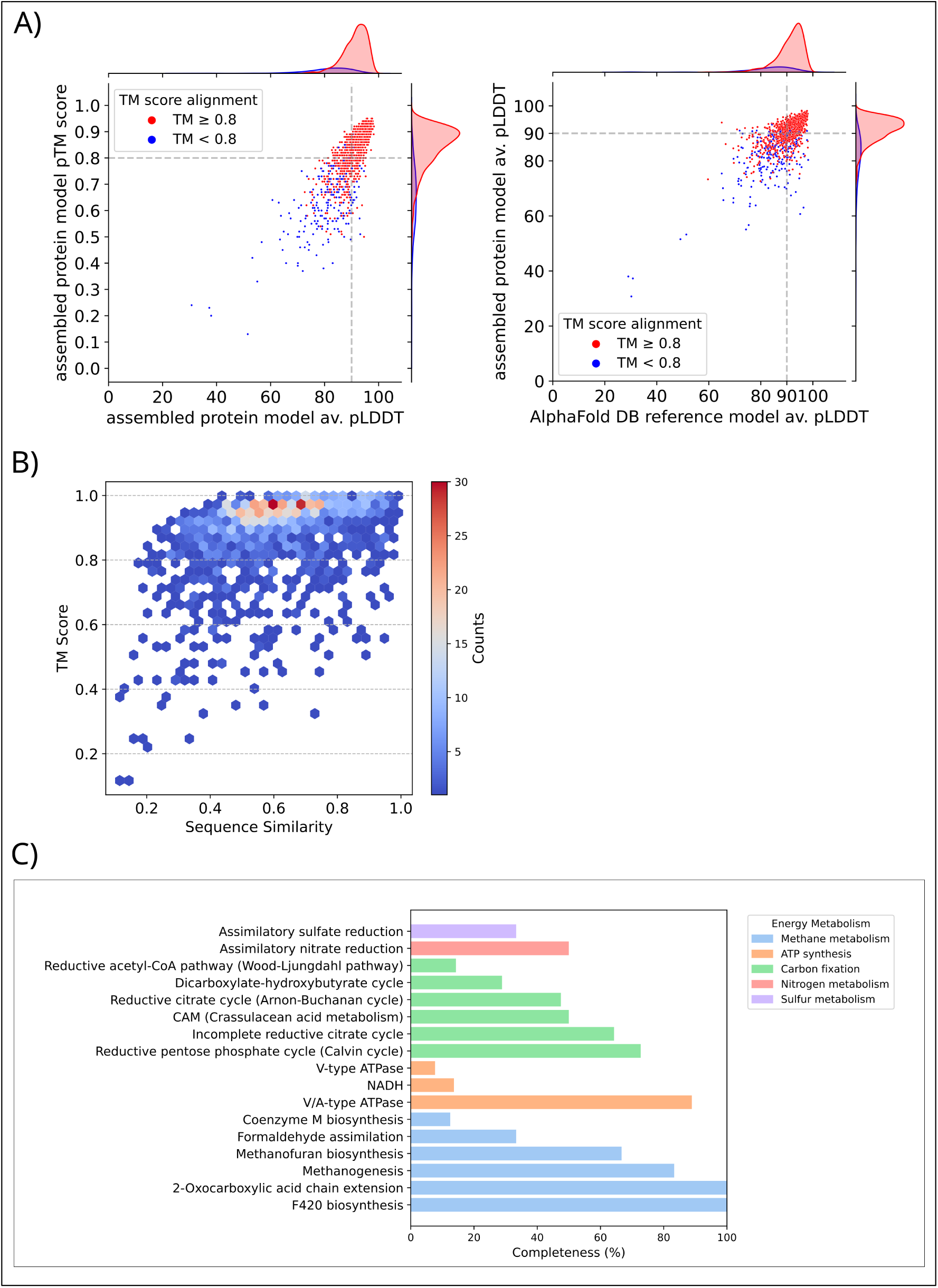
Structural and functional evaluation of ancient protein models. **Panel A:** Scatter plots of model confidence and alignment quality, colored by alignment TM score to the best AlphaFold db hit, with marginal density curves (left: pTM vs. average pLDDT of ancient models; right: average pLDDT of ancient models vs. average pLDDT of matched reference structures). Most models combine high confidence with strong structural similarity to modern homologues. **Panel B:** Hexbin plot of structural similarity (TM score) versus sequence similarity, colored by hit density. Most models retain high TM scores across sequence identities. **Panel C:** Completeness of KEGG modules by energy metabolism class, based on eggNOG-mapper annotation of the 1,299 representative proteins.

Figure **3**, **panel A** (left), shows a scatter plot of pTM versus average pLDDT for our ancient protein models, with marginal density curves on the top (pLDDT) and right (pTM) axes. A clear positive correlation is visible: models with low pLDDT generally also have low pTM. The density distributions highlight that models with both high pTM (*≥*0.8) and high pLDDT (*≥*90) predominantly correspond to high-confidence alignments to database entries, whereas those with pTM*<*0.8 tend to have lower pLDDT and align with lower confidence.

A similar pattern emerges when comparing the average pLDDT of the predicted ancient protein structure models with the average pLDDT of their best-matching AlphaFold db entry. Figure **3**, **panel A** (right), plots the reference average pLDDT on the x-axis against ancient-model average pLDDT on the y-axis, with points colored by alignment TM score (red for TM*≥*0.8). The marginal density curves show that most ancient models and their matched references have high average pLDDT values and high-confidence alignments, while lower average pLDDT in the ancient models corresponds to lower alignment scores.

Overall, these results demonstrate that ancient protein sequences reconstructed from highly fragmented aDNA can fold into structures closely resembling modern homologues. These results support the feasibility of retrieving functional proteins from ancient metagenomic datasets.

The most interesting proteins from an evolutionary perspective are proteins whose structures are conserved yet whose sequences diverge from known references [40]. To identify such proteins we compared the structural similarity (TM score) with sequence identity. Figure **3**, **panel B**, presents this comparison as a hexbin plot: TM scores on the y-axis versus sequence similarity on the x-axis, with bin color indicating hit density. While most protein structures achieve TM*≥*0.8 across a broad range of sequence identities (0.2–1.0), the highest density clusters near 0.6 sequence identity. Proteins of particular interest fall into the region with low sequence identity (0.2–0.4) but high TM scores (*≥*0.8), as these may represent ancient folds that have been maintained despite substantial primary-sequence divergence.

### 3.3 Pathway Analysis

To explore the functional roles of our ancient proteins beyond the basic “methan” annotation filter, we annotated all 1,299 representative sequences with eggNOG-mapper [29] (Section 3.6). This tool leverages both sequence- and profile-based searches of the eggNOG database [41] to infer orthologous relationships. The analysis resulted in 562 KEGG Orthology (KO) [30] identifiers. By filtering these by KO terms belonging to the “Energy Metabolism” category, we were left with 70 unique KO terms.

Figure **3**, **panel C**, shows the completeness of KEGG modules for each energy metabolism class. Methane metabolism modules are enriched: the *2-Oxocarboxylic acid chain extension* and *F420 biosynthesis* modules are 100% complete, while *Methanofuran biosynthesis* and *Methanogenesis* reach over 60% and 80% completeness, respectively. *Coenzyme M biosynthesis* and *Formaldehyde assimilation* were less than 40% complete. KEGG modules related to carbon fixation are also strongly represented, though none exceed 80% completeness. In contrast, the V/A-type ATPase module of the ATP synthesis pathway is nearly complete at *≈*90%. Overall, these results confirm an enrichment of methane metabolism among our selected proteins.

Since Figure **3**, **panels A** and **B**, report metrics for all filtered proteins prior to functional annotation, we summarise the model prediction and alignment statistics by pathway class in Table 4.3. It shows the median sequence similarity (as reported by Foldseek [5] alignments), median pLDDT for ancient models, median pLDDT for reference structures, and median TM score for each class. Across all pathways, confidence metrics are uniformly high: ancient-model average pLDDT medians range from 89.0 (ATP synthesis) to 95.3 (nitrogen metabolism), reference-model pLDDT from 90.7 to 97.0, and TM scores from 0.93 to 0.97. Sequence similarity varies by pathway, from a low median of 0.59 in ATP synthesis to a high of 0.79 in sulfur metabolism.

**Table 3:** Median sequence similarity, ancient-model pLDDT, database-model pLDDT, and TM score for each pathway class.

| Pathway Class | Median |  |  |  |
| --- | --- | --- | --- | --- |
|  | similarity | pLDDT<br>(ancient) | pLDDT<br>(database) | TM score |
| Methane metabolism | 0.62 | 94.04 | 95.54 | 0.95 |
| Carbon fixation | 0.69 | 94.34 | 95.66 | 0.97 |
| ATP synthesis | 0.59 | 89.00 | 90.74 | 0.93 |
| Nitrogen metabolism | 0.62 | 95.28 | 96.98 | 0.95 |
| Sulfur metabolism | 0.79 | 89.55 | 93.74 | 0.93 |

### 3.4 Case study: an archaeal protein involved in methanogenesis

To illustrate the results produced by our framework, we examined a candidate archaeal protein recovered from the approximately 2-million-year-old Kap København permafrost dataset. The predicted protein is highly similar to archaeal V/A-type ATP synthase subunit A, a catalytic component of the ATP synthase complex. Subunit A contains the nucleotide-binding sites required for ATP hydrolysis and synthesis and forms part of the catalytic headpiece of the enzyme. In methanogenic archaea, the V/A-type ATP synthase uses the ion gradient generated during metabolism to drive ATP synthesis, making this complex central to cellular energy conservation.

The ancient sequence corresponds to the CarpeDeam consensus of contig 69 B2 100 L0 KapK-12-1-36 0 carpe 000000047760 1 (ORF 253–1992, reverse strand). It is 1,740 nt long and can be aligned without gaps to a homologous coding sequence from the Euryarchaeota archaeon MAG MFD04408.bin.c.4 (OY969693.1:27356–29095). Across the coding sequence, we identified 118 substitutions relative to the modern comparator (Fig. **4**A; the complete column-by-column alignment is given in Supplementary Figure S1). These comprised 49 C*→*T/G*→*A substitutions, 31 reciprocal T*→*C/A*→*G substitutions, and 38 transversions; the full 12-change substitution spectrum is shown in Supplementary Figure S2. The whole-CDS divergence was 6.8%, with substitutions unevenly distributed along the sequence (Fig. **4**A). The coding impact of this divergence is limited: the 118 substitutions fall in 112 codons, only 29 of which alter the encoded residue, so 5.0% of the 579 residues differ between the two proteins (Supplementary Figure S3).

The most substitution-dense region occurs between nt 1576 and 1605, where eight substitutions affect six codons and four produce amino-acid changes (T527M, S532G, R533K and I535V) (Fig. **4**B). In contrast, the Walker A P-loop (GGFGTGKT; residues 227–234) is completely conserved. The corresponding 30-nt region contains a single synonymous C*→*T substitution, leaving the encoded motif unchanged (Fig. **4**C). Overall, 31 of the 118 nucleotide substitutions are non-synonymous, comprising 11 C*→*T/G*→*A transitions, 9 T*→*C/A*→*G transitions, and 11 transversions (Ti/Tv = 1.82; 95% Clopper–Pearson CI: 0.83–4.20) (Fig. **4**D). Deamination-driven damage would produce an excess of C*→*T/G*→*A over its reciprocal T*→*C/A*→*G, we observe a raw count ratio of 1.22 (95% CI: 0.46–3.34), consistent with no enrichment. Raw counts are not by themselves comparable, however, without accounting for the number of sites at which each base change could actually alter the encoded residue; normalising by these opportunity counts (586 and 581 sites, respectively) leaves an essentially unchanged 1.21-fold excess, which is not significant when tested against the null that the two classes split in proportion to their available sites (two-sided exact binomial *p* = 0.82; Supplementary Figure S4). We note that these counts give us limited power as a C*→*T/G*→*A excess below *∼*3-fold would not be detected at *α* = 0.05, so these data are compatible with the absence of post-mortem damage but do not completely reject a modest contribution.

Importantly, the substitution pattern of the fragments that have produced the contigs which have been realigned to it is consistent with post-mortem DNA damage rather than being driven primarily by a uniform excess of a particular substitution class. The 1,482 aDNA fragments contributing to the ancient consensus show the expected terminal enrichment of C*→*T substitutions at the 5*^′^*ends of fragments and the complementary G*→*A pattern at their 3*^′^* ends, reaching 34% and 36%, respectively, at the terminal position (Fig. **4**E). This provides independent support for the interpretation that the peptide sequence is of ancient origin.

**Figure 4:**
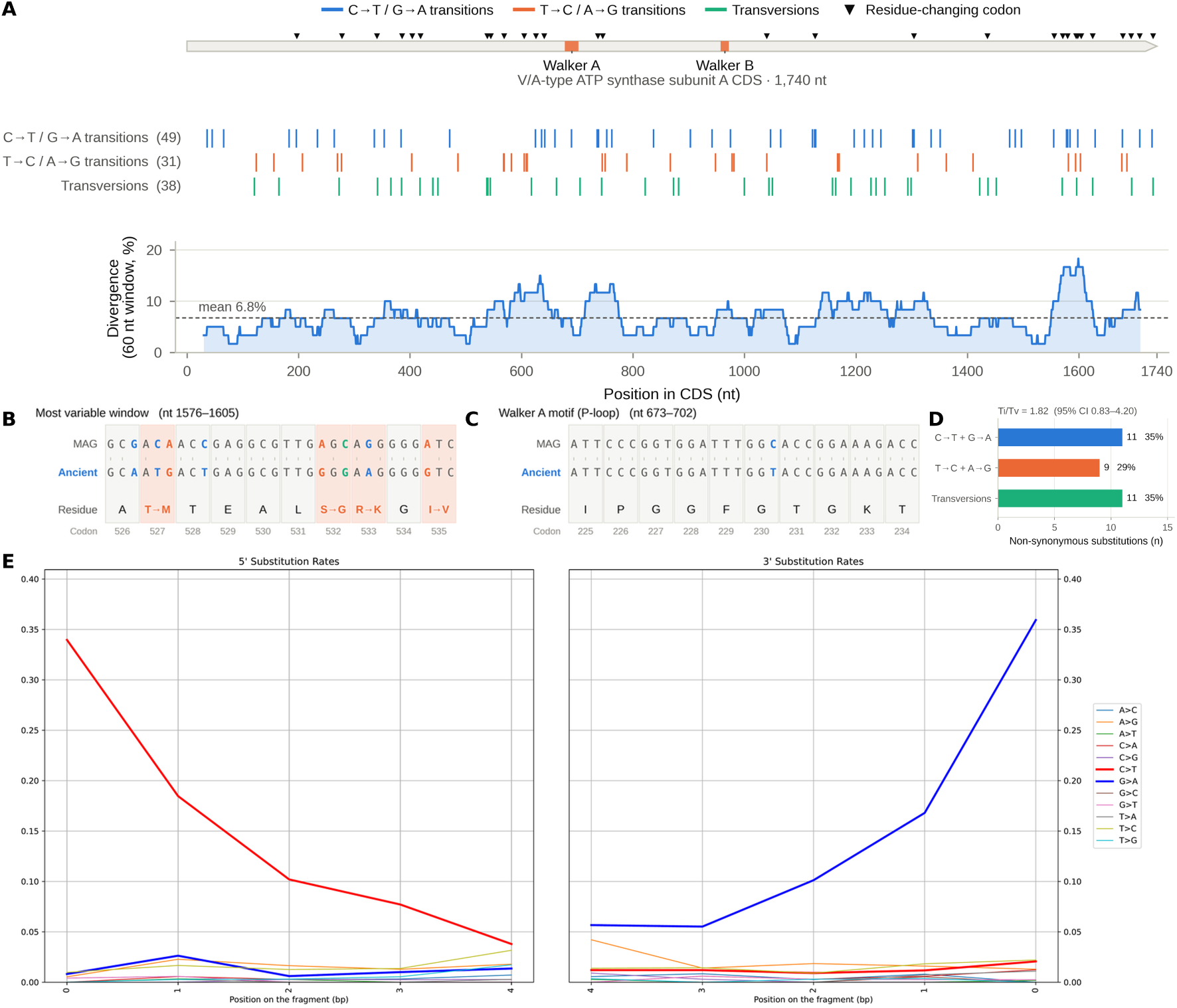
Ancient and modern archaeal ATP synthase subunit A sequences. The ancient sequence is the CarpeDeam consensus and the comparator is the corresponding CDS from the Euryarchaeota archaeon MAG MFD04408.bin.c.4. Substitutions are classified as C*→*T/G*→*A transitions (blue), reciprocal transitions (orange), or transversions (green). **(A)** Distribution of substitutions and nucleotide divergence along the CDS; Walker A and Walker B are indicated. **(B)** Most substitution-dense 30-nt window. **(C)** Walker A P-loop region. **(D)** Non-synonymous substitutions by substitution class. **(E)** Terminal nucleotide misincorporation frequencies in the 1,482 reads contributing to the ancient consensus.

We next asked whether the sequence similarity was accompanied by conservation of the protein’s predicted structure. The ancient sequence encodes a 579-residue protein, and its AlphaFold2 model is highly similar to an AlphaFold DB model of A-type ATP synthase subunit A from *Methanobacterium lacus* AL-21 (UniProt F0T967). Structural superposition using US-align [42] produced TM-scores of 0.986 and 0.979 when normalized by the ancient and reference lengths, respectively, with a Cα RMSD of 0.71 Å across 575 aligned residues. An independent PyMOL CEalign [43] analysis gave an RMSD of 0.86 Å across 568 residues. Thus, despite the age of the sequence, its predicted structure is essentially indistinguishable from that of a modern homolog (Fig. **5**A).

The structural similarity is also distributed across the protein rather than being driven by a small subset of residues. The median C*α* deviation is 0.50 Å, 99.1% of aligned residues fall within 2 Å, and the maximum deviation is 2.83 Å (Fig. **5**B). Notably, both the Walker A and Walker B motifs occur in regions of particularly low structural deviation, suggesting strong structural conservation at the two nucleotide-binding motifs (Fig. **5**B).

At the sequence level, the ancient protein is 95.0% identical to the modern MAG comparator and aligns without gaps over all 579 residues (Fig. **5**C). The 29 amino-acid differences are distributed throughout the sequence, while the Walker A (GGFGTGKT) and Walker B (ALMAD) motifs are identical in the two proteins. Only three of the 29 substitutions are both buried and non-conservative by BLOSUM62 (R209C, T527M and E347G), and none falls in either Walker motif (Supplementary Figure S5). The predicted structure is also well supported by the per-residue confidence scores: the mean pLDDT is 93.2, with a minimum of 52.1, and the majority of residues fall in the very-high-confidence range (Fig. **5**C; the model colored by pLDDT is shown in Supplementary Figure S6). Together, the sequence, damage, and structural evidence therefore support the interpretation that this ancient sequence derives from a highly conserved archaeal ATP synthase subunit A retaining both its characteristic catalytic motifs and overall protein fold.

**Figure 5:**
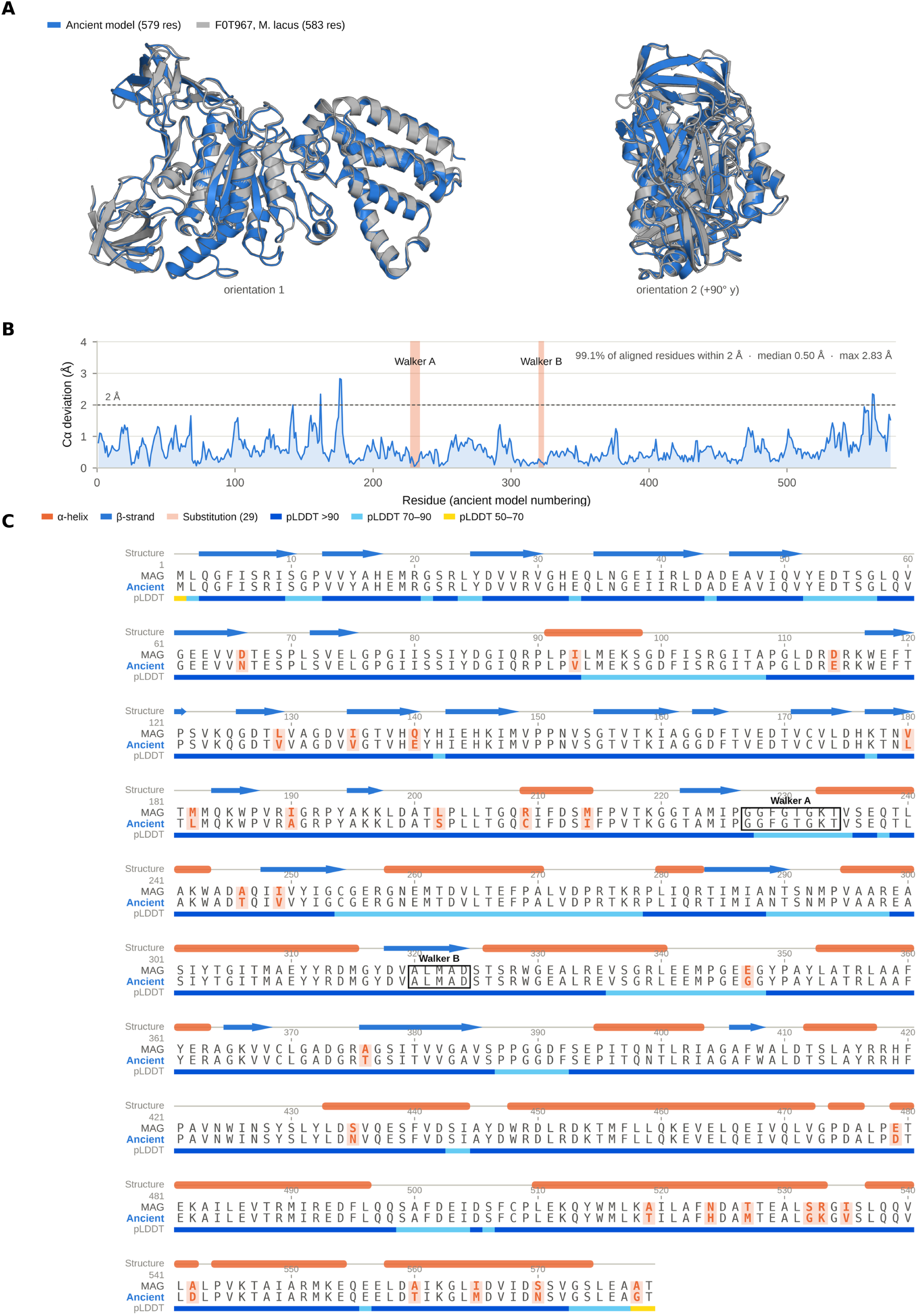
Structural and sequence conservation of the ancient archaeal ATP synthase subunit A. **(A)** Superposition of the ancient AlphaFold2 model and a reference AlphaFold model from *Methanobacterium lacus* AL-21. **(B)** Per-residue C*α* deviation in the structural superposition; Walker A and Walker B are indicated. **(C)** Alignment of the ancient sequence with the modern MAG comparator, with predicted secondary structure and per-residue pLDDT shown above the alignment.

## 4 Discussion

In this study, we demonstrate that protein-coding sequences can be recovered from ancient metagenomic datasets despite ultra-short fragment lengths and characteristic aDNA damage patterns. We investigated the applicability of three-dimensional structure prediction of ancient proteins for downstream functional analysis.

Instead of exhaustively folding the millions of sequences in our dataset, we adopted a depth-first workflow: we only kept predicted ORFs that match reference proteins presumably associated with methane metabolism. Then, we predicted their structures with AlphaFold 2 [1], and evaluated the resulting ancient protein structure models by aligning them to reference structures. This proof-of-concept shows that, using currently available tools, one can traverse from raw, damaged reads to full protein structures that align to modern homologs with high confidence. In the following, we evaluate potential technical refinements to the workflow and explore the biological implications of our results.

### 4.1 Assembly-to-Structure Pipeline Assessment

Our results confirm that the prediction of protein structures from ancient metagenomic datasets is feasible in practice. However, considerable challenges remain in the areas of assembly quality evaluation, damage authentication and computational resource requirements. Here, we highlight such challenges.

#### *De novo* assembly and ORF prediction are not error-free

Contigs produced by *de novo* assembly may incorporate incorrect bases due to aDNA damage [17] or form chimeric misassemblies that join fragments from unrelated genomic regions [15]. Translating nucleotide contigs into protein sequences and comparing predicted structures mitigates false positives: only one of the six possible reading frames encodes the correct protein, and amino acid sequences and three-dimensional folds are far more conserved than nucleotide sequences [44]. However, nonsynonymous substitutions or premature stop codons introduced by base substitutions [17, 10] or chimeric contigs [15] can reduce the spectrum of recoverable proteins. As a result, the proteins analysed may represent only a subset of all ancient proteins present in the data.

While our analyses focused on proteins that aligned to at least one target in the NCBI nr database, 76,482 ORFs did not match any entry. Several explanations may account for these “no-hit” sequences: (1) our chosen reference database does not capture full sequence diversity and does not contain the entirety of known sequences. A more comprehensive alternative would be the much larger MGnify database that contains over 2.4 billion sequences [45]. However, searching millions of contigs against it is computationally more intensive; (2) some sequences may be erroneous, arising from frame-shifted translations or chimeric contig assemblies; or (3) they may represent genuinely ancient, perhaps extinct, proteins. The latter scenario is particularly intriguing, but it can only be determined after systematically excluding possibilities (1) and (2).

#### aDNA authentication is a critical bottleneck in selecting protein sequences for structural modelling

Damage patterns are inferred by mapping sequencing reads against predicted ORFs from assembled contigs and analysing the resulting damage profiles [26]. However, read-mapping is prone to biases: ambiguous mappings can skew damage estimation and lead to incorrect assignments of reads to sequences (e.g. contigs). Such biases arise from the short fragment lengths and characteristic base substitutions of aDNA reads [46]. Furthermore, redundant reference sequences further amplify this problem. When multiple contigs cover the same genomic region, or when alternative ORFs are predicted from a single contig, reads may map equally well to any of these redundant sequences (in nucleotide space) [47]. In order to minimise false positives, most pipelines only retain the best-scoring alignment. This means that an ambiguous mapping read is only assigned to one of the redundant references [48, 12]. As a result, among a set of redundant reference sequences (e.g. contigs), only the one that has mapped sufficient reads to validate an aDNA damage signature is further analysed downstream. Other ORFs that potentially encode biologically interesting proteins, may be erroneously discarded simply because they lack enough mapped reads to demonstrate damage. Here, we discarded ORFs with fewer than 100 mapped reads, which eliminated over half of the candidate sequences. However, without this threshold, we would risk retaining damage patterns that are too weak to be reliable, which could lead to incorrect assumptions about the origin of the DNA. This threshold was also proposed in [47].

An alternative is to use all predicted ORFs for structural modelling, independently of their initial damage signatures, and postpone authentication until later stages. However, this strategy would substantially increase the computational burden, especially in terms of GPU resources required for protein folding [1, 28].

#### Predicted structures are not guaranteed to be accurate

Several metrics can validate protein structure models: the predicted Local Distance Difference Test (pLDDT) score, the predicted Template Modeling (pTM) score, and the TM-score from structural alignments to reference models [31]. The TM-score is particularly informative because it quantifies the structural similarity to a known structure. Therefore, a high TM-score supports the reliability of the predicted fold.

However, two key limitations remain. First, because our reference is also a computational model rather than an experimentally determined structure, its uncertainties affect the comparison. We addressed this by evaluating the pLDDT scores of both the ancient models and the references. Second, while high alignment scores confirm known folds, they offer limited insight into novel or mutated structures that are of greater interest.

In a recent study, Yeo *et al.* [49] applied a domain-centric analysis to over 800 million predicted structures from the ESMatlas [50] and the AlphaFold db [2] and found that re-predicting low-confidence models uncovered only a single novel fold, indicating near-saturation of known domain space. Extending this workflow to our pipeline could improve detection of ancient proteins with unknown structures and shed light on the evolutionary history of protein folds. If no new folds are uncovered, it would suggest that modern databases already capture most structural diversity. Alternatively, as Yeo *et al.* [49] suggest, we may have reached the current limits of structure prediction models. However, implementing this approach would demand substantial computational resources. First to fold the millions of ORFs extracted from our contigs, and then to search them against the database of over 800 million predicted structures.

### 4.2 Assessment of Results from a Biochemical Perspective

Our findings revealed ancient protein structures that describe the methanogenesis pathway almost completely. These results provide a framework for investigating diverse topics of methane-related research.

Methane is a potent greenhouse gas. Understanding its production pathways has significant implications for climate research [51, 52]. For instance, it is estimated that around 2.5 million years ago, Greenland’s climate resembled the conditions forecasted under future warming scenarios [9]. The accelerated thawing of present-day permafrost exposes previously frozen anaerobic wetlands, creating environments in which methanogenic microbes can thrive and potentially significantly increasing global methane emissions [52, 53, 54].

Several studies have examined the genetic diversity of methanogens and microbial communities from past warm periods using ancient samples containing either degraded DNA [55, 10, 56] or intact, dormant organisms [57, 58, 59]. Similar to our workflow Perfumo *et al.* [55] and Orsi *et al.* [56] predicted protein sequences from *de novo* assembled contigs for homology searches and sequence-based annotation. Extending these methods with protein structure modelling has great potential to provide more refined insights: it enables the validation of assembled proteins and may reveal homologues with low sequence similarity (*<* 20%) due to the highly conserved nature of protein folds [44], as well as allowing the analysis of functional efficiency [60]. Until recently, such analyses required crystallographic structures from living samples [61, 62], which are costly and labour-intensive to obtain [57].

Overall, integrating structure-based approaches into ancient metagenomic workflows opens new possibilities for studying methanogenic microbes in the context of climate change and investigating the evolutionary origins and diversification of methanogenesis. The latter complements the ongoing efforts to map the full diversity of methanogens and their metabolic pathways [63, 64, 65, 66, 67].

### 4.3 Limitations of the structure-based approach

The archaeal ATP synthase subunit A provides a more detailed example of what can be learned once individual ancient proteins are examined beyond their initial database annotation. The sequence is supported by characteristic terminal aDNA damage in the reads contributing to its reconstruction, while the predicted protein retains the conserved Walker A and Walker B motifs and adopts a structure that is highly similar to modern homologues. This combination of independent evidence is important because sequence similarity alone cannot distinguish a genuinely ancient protein from a modern contaminant that happens to match the same reference. At the same time, the close structural conservation of this protein illustrates a limitation of structure-based analyses: recovering a structure that is nearly identical to a modern homologue provides strong support for its identity and fold, but offers limited evidence for ancient biochemical innovation. More divergent proteins, particularly those with weak sequence similarity but conserved structural features, will therefore be of greater interest in future applications of the framework.

## 5 Declarations

### Code Availability

The code for the analysis presented in this study are accessible under https://github.com/LouisPwr/AncientReconstructedProteins. Data supporting the findings of this study are deposited in Zenodo (https://doi.org/10.5281/zenodo.22027760).

### Hardware

Computationally intensive analyses, such as the large-scale database searches, were performed on servers with 5th Gen Intel^®^ Xeon^®^ Scalable Processors—either model 6548N or 8562Y+—each featuring 32 cores, 64 threads, a base frequency of 2.8 GHz, and a maximum turbo frequency of 4.1 GHz.

Protein structure prediction was performed on servers equipped with either NVIDIA H100 NVL GPUs (96 GB memory, CUDA 12.2) or NVIDIA L40S GPUs (46 GB memory, CUDA 12.2), depending on availability.

### Author Contributions

G.R. conceived the project. L.K. further developed the concept and designed and implemented the workflow. G.R. and L.K. performed the analysis and wrote the manuscript. P.W.S. provided help with the server infrastructure.

## Acknowledgements

L.K. acknowledges the use of OpenAI’s ChatGPT-4o and DeepL Write for proofreading (spelling, grammar, punctuation). L.K. was supported by a PhD stipend from Novo Nordisk Data Science Investigator grant (NNF20OC0062491). G.R. acknowledges the use for producing the code to produce Figure 4 and 5. The authors take full responsibility for any errors due to this process.

