## Supplementary Material for "Ancient Reconstructed Proteins: A Framework for Resurrecting Protein Structures from Million-Year-Old Metagenomes"

### Contents

S1 Case study: additional analyses of the archaeal V/A-type ATP synthase subunit A 3

#### S1 Case study: additional analyses of the archaeal V/A-type ATP synthase subunit A

The figures in this section accompany the case study presented in the main text (Figures 4 and 5). They report the analyses of the same two sequences — the `CarpeDeam` consensus of contig 69_B2_100_L0_KapK-12-1-36_0_carpe_000000047760_1 (ORF 253–1992, reverse strand) and the corresponding coding sequence of the Euryarchaeota archaeon MAG MFD04408.bin.c.4 (OY969693.1: 27356–29095) — that are not shown in the main figures. Throughout, substitutions are polarised comparator $\rightarrow$ ancient and grouped into the same three classes used in the main text: C$\rightarrow$T / G$\rightarrow$A transitions (blue), which are the two strand orientations of the post-mortem cytosine deamination signature; the reciprocal T$\rightarrow$C / A$\rightarrow$G transitions (orange); and transversions (green).

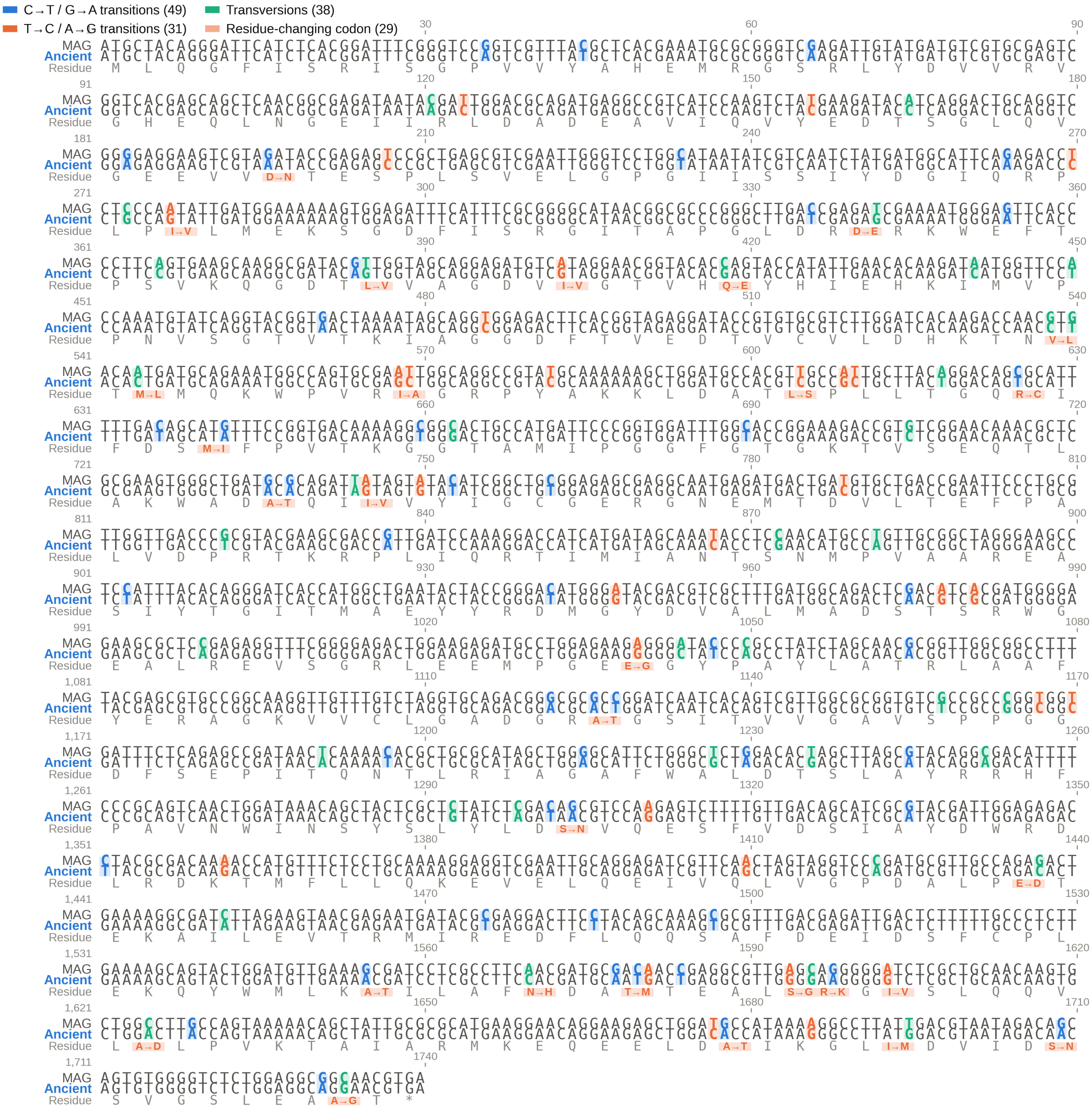


**Figure S1**: **Complete nucleotide alignment of the ancient CDS against its modern comparator.** The archival companion to Figure 4A of the main text: all 1,740 aligned columns are shown in 20 blocks of 90 nt (30 codons per block, so the reading frame stays in register), comparator ("MAG") above and ancient below, with a ruler every 30 nt and the absolute coordinate of the first base at the start of each block. Substituted columns are shaded and bold and coloured by class: C→T / G→A transitions (blue, $n = 49$), T→C / A→G transitions (orange, $n = 31$) and transversions (green, $n = 38$). The translated residue is carried on a third row beneath each codon, so synonymous changes are visible as such: where the substitution alters the residue the codon is shaded orange and the change written as X→Y, and where it does not the single shared residue is shown in grey. 29 of the 112 affected codons change the encoded residue.

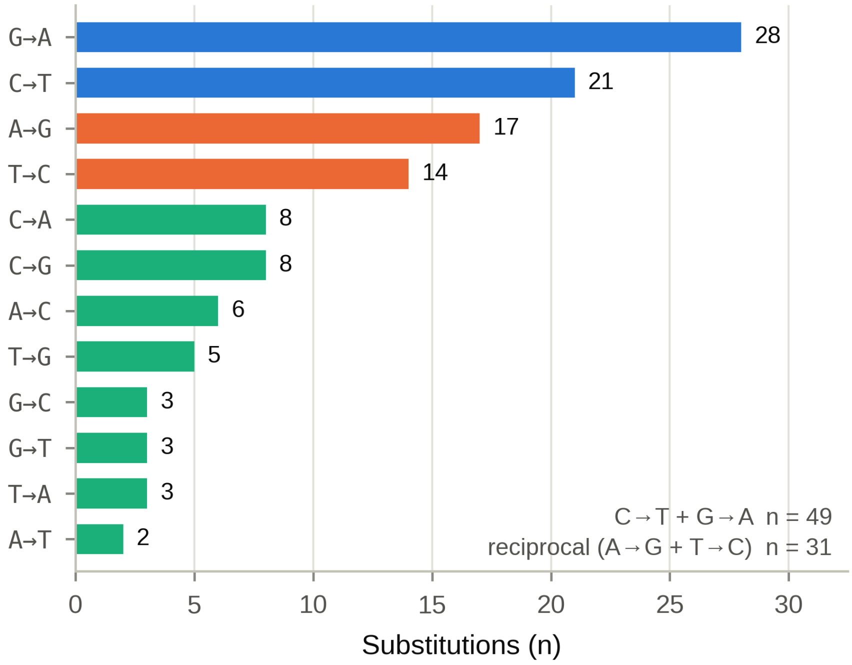


**Figure S2**: **Substitution spectrum of the ancient coding sequence against its modern MAG comparator.** All 12 base changes, polarised comparator → ancient, ordered by count and coloured by class. The deamination-like changes total 49 (G→A 28, C→T 21) against 31 for their reciprocal transitions (A→G 17, T→C 14), out of 118 substitutions in 1,740 nt. C→T and G→A are the two strand orientations of the same post-mortem cytosine deamination event, which is why they are counted together and separated from the other transitions.

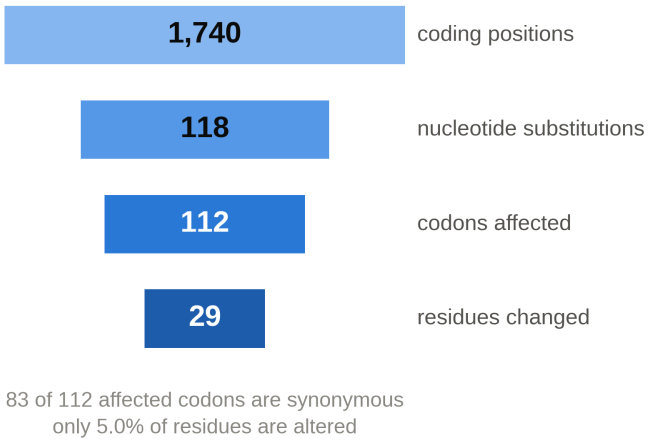


**Figure S3**: **Coding impact of the nucleotide divergence, as a funnel from coding positions to altered residues.** Of 1,740 coding positions, 118 carry a nucleotide substitution; these fall in 112 codons; and only 29 of those codons change the encoded residue. 83 of the 112 affected codons are therefore synonymous, and 5.0% of the 579 residues are altered.

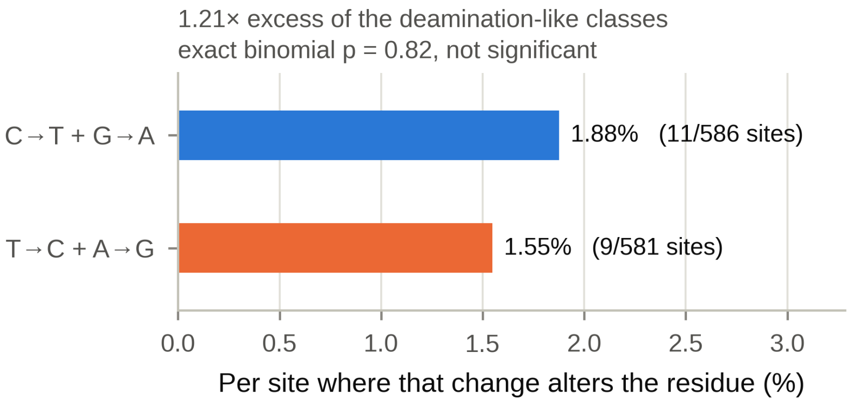


**Figure S4**: **Per-site rates of the two transition classes among the non-synonymous substitutions.** Each class's count is divided by the number of sites in the comparator coding sequence where that particular base change would actually alter the residue: 11 of 586 such sites (1.88%) for C→T + G→A, and 9 of 581 (1.55%) for T→C + A→G. The correction is needed because the raw counts shown in Figure 4D of the main text are not comparable without knowing how many sites each class had to work with. Here the two opportunity counts are close to parity, but that is a result rather than an assumption. The residual 1.21-fold excess of the deamination-like classes is not significant: exact two-sided binomial $p = 0.82$, tested against the null that the two classes split in proportion to their available sites rather than against 0.5. All binomial tests and exact confidence intervals were computed from scratch.

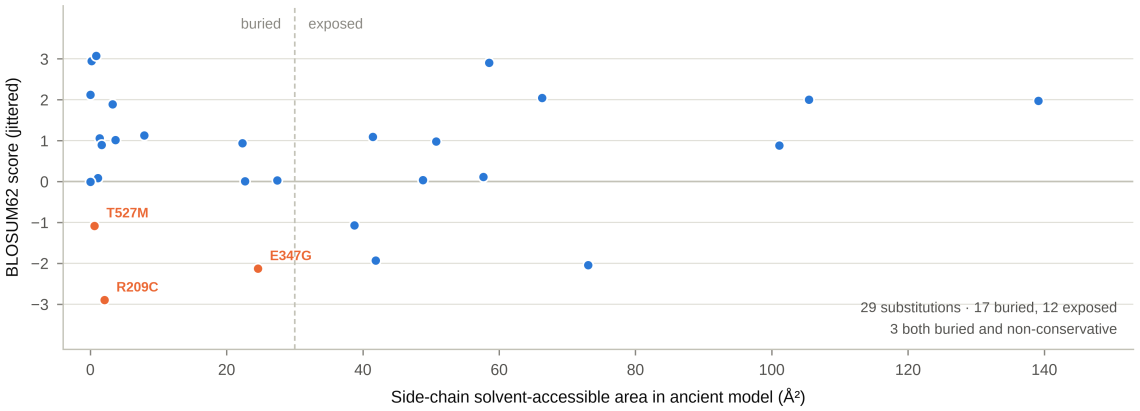


**Figure S5**: **Character of the 29 amino-acid substitutions between the ancient subunit A and its modern MAG comparator.** BLOSUM62 exchange score against the side-chain solvent-accessible area of that residue in the ancient ColabFold model. The vertical dashed line at 30 $Å^2$ separates buried from exposed side chains: 17 substitutions are buried and 12 exposed. Only three are both buried and non-conservative (BLOSUM62 $< 0$) and are labelled in orange — R209C, T527M and E347G; the remaining 26 are drawn in blue. No substitution falls in either Walker motif. BLOSUM62 scores are integers; points are jittered vertically only to separate coincident markers.

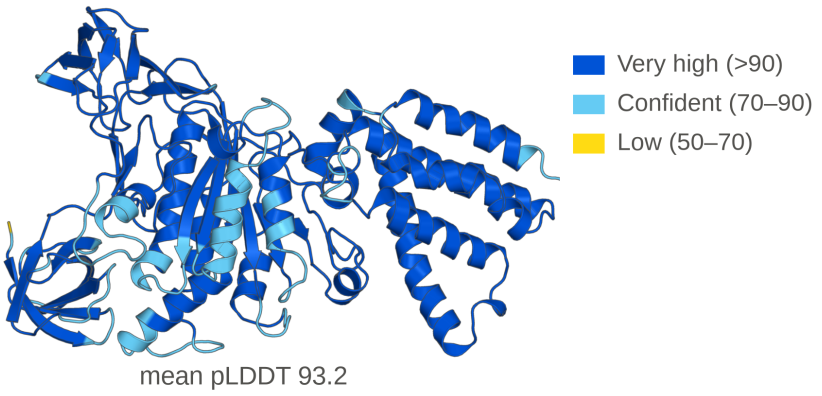


**Figure S6**: **The ancient subunit A model coloured by per-residue prediction confidence (pLDDT).** Colours follow the AlphaFold convention: dark blue, very high ($> 90$); light blue, confident (70–90); yellow, low (50–70). The “very low” band ($< 50$) does not occur — the model’s minimum pLDDT is 52.1. The model is the ColabFold AlphaFold2-ptm prediction of the `CarpeDeam` consensus (579 residues), shown in the same orientation as the left-hand view of Figure 5A of the main text. Mean pLDDT 93.2.
